# The effect of known mutator alleles in a cluster of clinical *Saccharomyces cerevisiae* strains

**DOI:** 10.1101/090498

**Authors:** Daniel A. Skelly, Paul M. Magwene, Brianna Meeks, Helen A. Murphy

## Abstract

Natural selection has the potential to act on all phenotypes, including genomic mutation rate. Classic evolutionary theory predicts that in asexual populations, mutator alleles, which cause high mutation rates, can fix due to linkage with beneficial mutations. This phenomenon has been demonstrated experimentally and may explain the frequency of mutators found in bacterial pathogens. In contrast, in sexual populations, recombination decouples mutator alleles from beneficial mutations, preventing mutator fixation. In the facultatively sexual yeast *Saccharomyces cerevisiae*, segregating alleles of *MLH1* and *PMS1* have been shown to be incompatible, causing a high mutation rate when combined. These alleles had never been found together naturally, but were recently discovered in a cluster of clinical isolates. Here we report that the incompatible mutator allele combination only marginally elevates mutation rate in these clinical strains. Genomic and phylogenetic analyses provide no evidence of a historically elevated mutation rate. We conclude that the effect of the mutator alleles is dampened by background genetic modifiers. Thus, the relationship between mutation rate and microbial pathogenicity may be more complex than once thought. Our findings provide rare observational evidence that supports evolutionary theory suggesting that sexual organisms are unlikely to harbor alleles that increase their genomic mutation rate.

## Introduction

Mutation rates, like most organismal phenotypes, are subject to natural selection[1]. Alleles that increase the genomic mutation rate can be subject to direct selection but also to indirect selection on the fitness effects of mutations at other loci. A rich body of evolutionary theory predicts that in asexual populations, mutator alleles with little or no direct fitness effect can rise to fixation when they are linked to rare beneficial mutations. The probability of mutator hitchhiking is related to the supply and magnitude of such mutations, as well as the population size [2-6]. Experimental studies with *Escherichia coli* [7-12] and *Saccharomyces cerevisiae* [13-15] have provided evidence for this phenomenon, and clinical isolates of asexual pathogenic microbes have been shown to contain elevated frequencies of mutator strains [16-20], suggesting that mutator alleles have a propensity to rise in frequency during invasion of a new ecological niche. In contrast to asexual populations, mutator hitchhiking is not predicted to occur in sexual organisms. According to simulations and theoretical models, sex and recombination will erode linkage, separating beneficial mutations from the mutator alleles that caused them, and preventing an increase in frequency of the mutators [5, 21-25]. This prediction has garnered experimental support in *Saccharomyces cerevisiae* [26]. However, some population genetic models have shown that under certain restrictive circumstances, fluctuating environments could select for mutator alleles in sexual populations[27-29].

Sexual microbes in the real world are mostly eukaryotes with facultatively sexual life cycles that undergo infrequent sex between periods of clonal growth [30]. The degree of linkage within the genome, as well as the strength and direction of selection and environmental change, are usually unknown. Thus, it is unclear whether natural populations of sexual microbes are expected to contain considerable frequencies of mutator strains, and to our knowledge none have been reported.

The budding yeast *Saccharomyces cerevisiae* has been collected from locations and ecological settings around the world [31-34]. Its lifecycle can include asexual growth in haploid and diploid forms, as well as both outcrossing and extreme inbreeding (mating-type switching and intra-ascus mating) [35]. Outcrossing rates are inferred to be extremely low in natural populations with most growth occurring clonally in the diploid phase [36, 37]. *S. cerevisiae* is also known to be an opportunistic pathogen [38, 39]. Pathogenic isolates have been shown to have high levels of heterozygosity [40], suggesting that sexual outcrossing events prior to colonization of a new ecological niche may be associated with opportunistic pathogenicity. The global population of *S. cerevisiae* contains two incompatible alleles (which we will refer to as the “mutator” allele combination) in the genes *MLH1* and *PMS1* [41] that, when engineered together, increase the mutation rate 20-400X above wild type [42]. Until recently, no natural isolates had been found carrying both alleles, in line with theoretical expectations of the absence of mutators in sexual populations. However, strains recently sequenced by Strope et al. (2015) include a cluster of clinical isolates that carry both mutator alleles. This cluster of strains could represent a rare case of a facultatively sexual microbe with a naturally occurring mutator phenotype, and one that may be associated with invasion into a new ecological niche (the human body).

Here we report direct estimates of the genomic mutation rates in the clinical background containing the two incompatible alleles. Furthermore, we investigate the phylogenetic history of the mutator alleles and examine levels of genomic variation in both mutator and non-mutator genetic backgrounds. We find that these isolates have a mildly elevated mutation rate (~4 times greater than closely related non-mutator strains), significantly lower than previously reported for incompatible combinations of *MLH1* and *PMS1* alleles assayed in a single laboratory genetic background [42]. Genomic analysis of mutator and non-mutator strains provided no evidence of a historically elevated mutation rate. We conclude that the mutational effect of the incompatible mismatch repair alleles is dampened by background genetic modifiers, and discuss possible explanations for our observations.

## Materials and Methods

### Strains and Media

Fifteen strains were chosen from the 100 Genomes Yeast panel [34]: four that contained both of the alleles necessary to confer a mutator phenotype (MLH1^D761^ and PMS1^K818^), six that contained only the PMS1^K818^ allele, and five that contained only the MLH1^D761^ allele. Two MLH1^D761^-PMS1^K818^ strains that were reported to have a mutation rate ~100X wild type [41], EAY1370 and EAY1363 (generously provided by Eric Alani), were used as a control. All strains assayed were heterothallic haploid MATa (i.e., derived from the original diploid background). In order for the control strains to be used in the fluctuation assays, the *URA3* gene was restored via lithium acetate transformation [43] using strain YJM128 as the PCR template. Strains were Sanger sequenced at the *PMS1* and *MLH1* loci with the following primers to verify allele status: MLH1for-

GCAGGTGAGATCATAATATCCC; MLH1rev-
GGGCATACACTTTCAAATGAAACAC; PMS1for-
CAGATAAACGATATAGATGTTCATCG; PMS1rev-
CCTTCGAAAATGAGCTCCAATCA). Strains are listed in Table 1.

**Table 1:** Strains used in this study.

| Strain | Diploid Parent | Genetic Background | Genotype |
| --- | --- | --- | --- |
| YJM1775 | YJM320 | 91-190 | hoΔ::loxP, Mat a, PMS1<sup>K818</sup> |
| YJM1639 | YJM451 | B70302(b) | hoΔ::loxP, Mat a, PMS1<sup>K818</sup> |
| YJM1751 | YJM541 | YJM522 | hoΔ::loxP, Mat a, MLH1<sup>D761</sup>, PMS1<sup>K818</sup> |
| YJM1753 | YJM554 | YJM521 | hoΔ::loxP, Mat a, MLH1<sup>D761</sup>, PMS1<sup>K818</sup> |
| YJM1785 | YJM555 | YJM523 | hoΔ::loxP, Mat a, MLH1<sup>D761</sup>, PMS1<sup>K818</sup> |
| YJM1787 | YJM681 | YJM653 | hoΔ::loxP, Mat a, PMS1<sup>K818</sup> |
| YJM1789 | YJM682 | R87-91 | hoΔ::loxP, Mat a, MLH1<sup>D761</sup> |
| YJM1851 | YJM1083 | NRRL Y-10988 | hoΔ::loxP, Mat a, PMS1<sup>K818</sup> |
| YJM1747 | YJM1133 | MMRL 125 | hoΔ::loxP, Mat a, MLH1<sup>D761</sup>, PMS1<sup>K818</sup> |
| YJM1805 | YJM1199 | 1566 (UCD-FST 08-199) | hoΔ::loxP, Mat a, MLH1<sup>D761</sup> |
| YJM1673 | YJM1332 | M1-2 | hoΔ::loxP, Mat a, MLH1<sup>D761</sup> |
| YJM1876 | YJM1433 | Yllc17_E5 | hoΔ::loxP, His<sup>+</sup>, Mat a, PMS1<sup>K818</sup> |
| YJM1707 | YJM1444 | UWOPS87-2421 | hoΔ::loxP, Mat a, PMS1<sup>K818</sup> |
| YJM1825 | YJM1549 | DBVPG6040 | hoΔ::loxP, Mat a, MLH1<sup>D761</sup> |
| HMY257 |  | S288c (EAY1370) | hoΔ, Mat alpha, leu2, trp1, lys2::insE-A14, PMS1<sup>K818</sup>, MLH1<sup>D761</sup> |
| HMY258 |  | SK1 (EAY1363) | hoΔ, Mat a, lys2::insE-A14, leu2, his4xB, ade2, MLH1<sup>D761</sup>, PMS1<sup>K818</sup> |

Strains were grown in YPD [44]; spontaneous *ura-* mutants were selected on synthetic complete agar plates [44] supplemented with 1mg/ml 5-fluoro-orotic acid (5FOA) and 60mg/ml uracil. Cultures were also treated with deflocculation buffer (20 mM sodium citrate, 5 mM EDTA), as strains YJM1639, YJM1851, YJM1747, and YJM1707 flocculated.

### Fluctuation Assays

The modified Jones protocol was used to estimate mutation rates [45]; following the procedure of [26], spontaneous *ura-* mutants were assayed. All strains were grown overnight in YPD from freezer stocks; a new tube of 10ml of YPD was inoculated with ~1000 cells and grown for 48 hrs. Five replicate cultures of 30ml YPD were then inoculated with ~100-500 cells and grown for 48 hrs. The cultures were centrifuged, resuspended in 5ml deflocculation buffer, washed, and resuspended in 5ml or 10ml of water. For each replicate: (1) 100ul of an appropriate dilution was plated on YPD to estimate the population size; (2) 300ul was plated on 5FOA plates to select for spontaneous *ura-* mutants; (3) for control strains, 10ul + 90ul water was also plated on 5FOA plates. Mutation rates were estimated using “Mutation Rate Calculator” software (provided by P.D. Sniegowski and P. J. Gerrish). This assay was replicated four times, with at least one control mutator strain included in two of the replicates.

### Statistical tests for mutation rate variation

We used R version 3.2.2 [46] and the lme4 package [47] to implement mixed models to test for differences in mutation rate. For all models we used the logarithm of mutation rate as the response variable. We modeled assay as a random effect since particular levels are not of primary interest. We used likelihood ratio tests to test for an assay effect in models containing a random strain effect and fixed effect for mutator alleles. To test for differences between groups of strains (engineered mutators, natural mutators, and non-mutators), we included a random strain effect and a fixed effect for group differences. Coefficient estimates from these linear models quantified the fold differences between groups.

### Haplotype analyses

For each of the 100 yeast genomes studied by Strope et al. (2015) (except strain M22 which has a large amount of incomplete sequence), we extracted at least 10kb of DNA sequence surrounding the mutator alleles, amino acid 761 of *MLH1* and 818 of *PMS1*. We used a method developed by Gabriel et al. (2002)[48] and implemented in the program Haploview [49] to partition the sequences into blocks that did not show strong evidence of historical recombination. We examined the sequence block containing the site of the mutator allele substitution in each gene. We constructed haplotype networks using PopART (http://popart.otago.ac.nz) with the median joining network option [50].

### DNA sequence variation analyses

Complete genomes of the 15 strains whose mutation rates were measured were obtained from Strope et al. (2015). The genome of *S. paradoxus* was obtained from [51]. These 16 complete genomes were aligned with mugsy v1r2.2 [52] using default options. MAF files were manipulated using maftools v0.1 [53]. To clarify comparisons, only alignment blocks where sequences from all 16 taxa aligned were considered. The majority of sequence fell into this subset of the full alignment – for the 12.16Mb S288c genome, 11.7Mb (96%) was present in the complete alignment and 10.94Mb (90%) was present in alignment blocks containing sequence from all 16 taxa. In order to obtain a reliable set of polymorphic sites, the following criteria were used to filter further: (1) ends of alignment blocks were trimmed if the sequence of any species consisted of only gaps, (2) only biallelic single nucleotide variants were considered, not indels or multi-allelic sites, (3) for a particular site, any strains with a gap or N at that site were ignored.

## Results and Discussion

### Four strains derived from natural isolates carry incompatible mismatch repair alleles

Strope et al. (2015) sequenced the genomes of 93 *S. cerevisiae* strains derived from isolates collected globally with a particular focus on clinical isolates. These strains were haploid or homozygous diploid segregants of the isolates themselves. Surprisingly, four of the sequenced strains carried incompatible alleles in the mismatch repair genes *MLH1* (G761D) and *PMS1* (R818K) that are thought to confer a dramatic increase in mutation rate when they co-occur [41]. While both alleles are known to be segregating in natural populations (24% and 12%, for *MLH1^D761^* and *PMS1^K818^*, respectively), it was previously postulated that the alleles would not be found together in nature because the combination would be selected against due to the accumulation of deleterious mutations [41]. Indeed, a previous investigation into a diverse panel of yeast isolates uncovered no strains with the combination [42]. The four strains recently discovered to be carrying this “mutator” allele combination are all clinical isolates classified as admixed “mosaics” which do not show pure ancestry from a single *S. cerevisiae* population [34]. The isolates from which these strains derived were collected from geographically disparate locations in California (n=3) and North Carolina (n=1).

To understand the history of mutator alleles at *MLH1* and *PMS1*, we analyzed the haplotype structure of regions surrounding these loci in the Yeast 100 Genomes panel [34]. We partitioned this sequence into blocks showing little evidence of historical recombination, and found that each derived allele appears to have arisen once (on a single haplotype; Figure 1) and spread widely within *S. cerevisiae*, as evidenced by its presence among diverse wine/European and mosaic strains. This observation is expected due to the small or nonexistent fitness effect of each mutator allele when found with a wild-type allele at the opposing locus [54].

**Figure 1:**
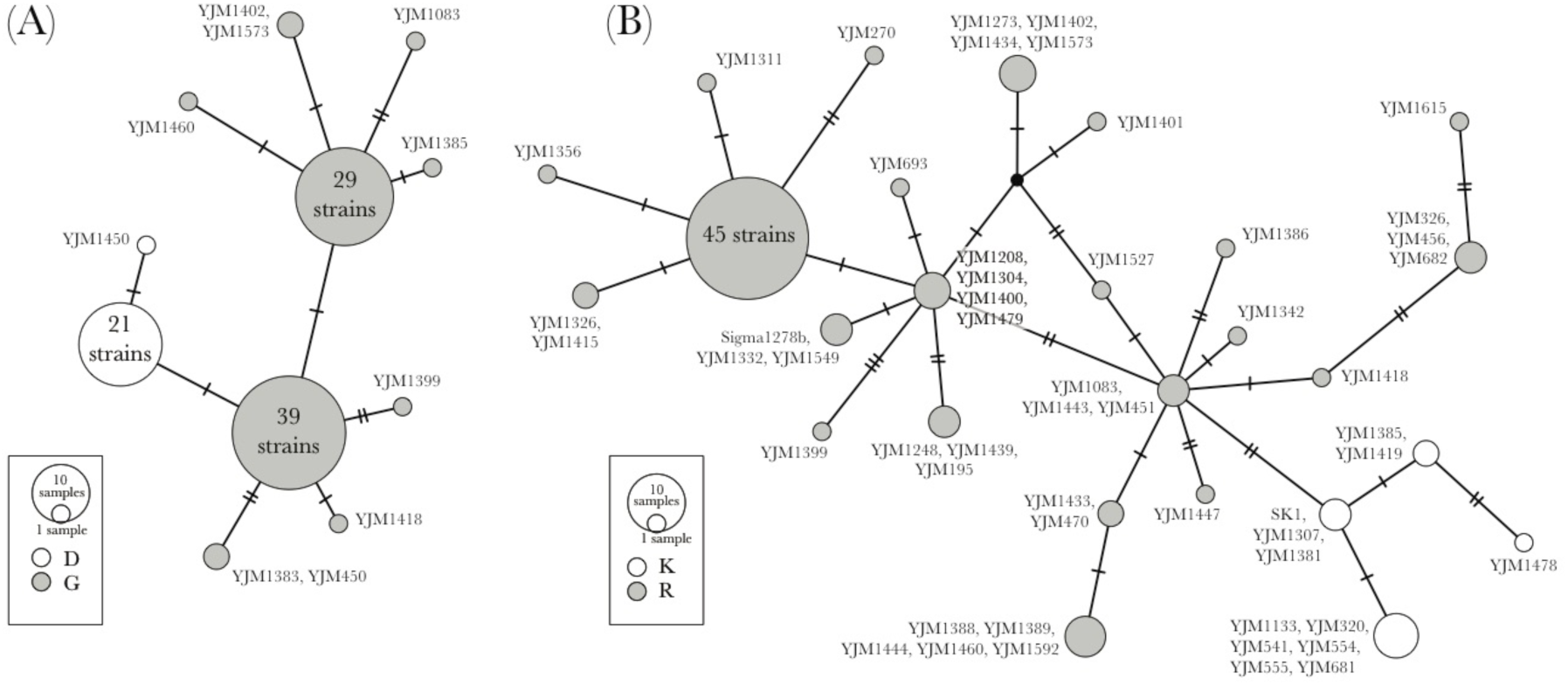
Haplotype networks constructed using variation surrounding mutator alleles. (A) *MLH1* and (B) *PMS1* networks. In both networks, white circles indicate haplotypes carrying the derived allele (the mutator combination consists of the derived allele at both loci). Both networks are derived from sequence data of 99 strains, including all strains assayed in this study (Methods).

### Genomic mutation rate is only slightly elevated in natural mutator strains

We assayed mutation rate in the four strains carrying the two incompatible mismatch repair alleles, 11 related strains carrying compatible combinations of alleles, and two laboratory strains engineered to carry the incompatible alleles and previously shown to exhibit high mutation rate (control strains). We refer to these strains as *natural mutator* strains, *non-mutator* strains, and *engineered mutator* strains, respectively. Although the strains we studied were derived from diploid isolates that could be heterozygous for the incompatible alleles, these strains have direct relevance to *S. cerevisiae* population biology: a single round of sporulation followed by intra-ascus mating or mother-daughter mating could produce a diploid yeast cell homozygous for both mutator alleles. We measured mutation rate multiple times (median 4, range 3-5) for each strain, except the two previously assayed engineered mutator strains (*n* = 1 or 2) and a single non-mutator strain (PMY1798; *n* = 1).

Given the large number of mutation rate measurements conducted in this study, we carried out the mutation rate assay in multiple batches and attempted to include all strains in each batch. Experimental variation led to significant variation in mutation rate estimates between batches (χ^2^_1_ = 9.6, *p* = 0.002). Thus, we used mixed models including assay as a random effect to test for differences between strains. As expected, the engineered mutator strains, previously demonstrated to exhibit a high mutation rate, had consistently and significantly higher mutation rate estimates than any other strains (Fig. 2; 40 times greater than the average non-engineered strains; χ^2^_1_ = 18.7, *p* = 1.56×10^-5^).

**Figure 2:**
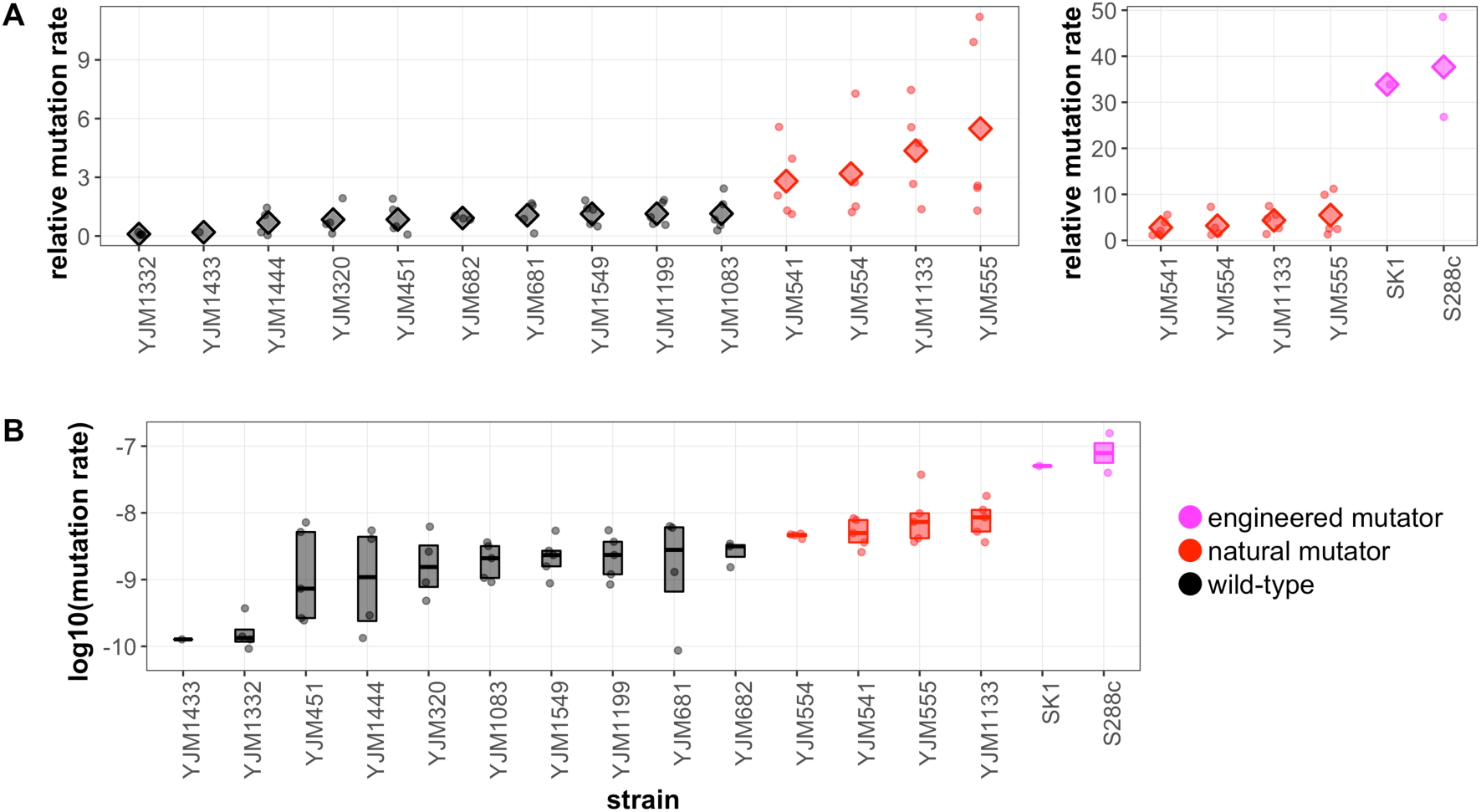
Mutation rates of strains carrying mutator and non-mutator allele combinations. (A) For each fluctuation assay, strains with only one of the mutator alleles were averaged to provide the baseline mutation rate; the mutation rate for each strain was divided by this baseline. Circles show data from individual assays and diamonds indicate average for the strain over all the assays. Left panel shows strains sequenced by Strope et al. (2015) and assayed in this study. Right panel compares the mutation rates of engineered mutator strains from Heck et al. (2006) to 100 genomes natural mutator strains. Note the difference in y-axis scale between the panels. (B) Estimates of mutation rate by strain. Dots indicate measurements from individual assays. Boxes show 25%, 50%, and 75% quantiles. In both panels, strain labels indicate genetic background to facilitate comparison to genomes sequenced by Strope et al. (2015).

Surprisingly, the four natural mutator strains had only slightly higher mutation rates than the non-mutator strains (Fig. 2; 5.6 times greater than non-mutator strains; χ^2^_1_ = 29.5, *p* = 5.65×10^-8^). Comparing the mutation rate estimates for these four strains to their two closest non-mutator relatives among the strains we assayed (YJM1775 and YJM1787), the difference was even more subtle (4.3 times greater; χ^2^_1_ = 11.2, *p* = 8.2×10^-4^). The specific alleles found in the clinical isolates show a significant increase in mutation rate when engineered into a lab background (Eric Alani, personal communication).

### Patterns of genetic variation are similar in mutator and non-mutator strains

Although the mutator allele combination did not appear to give rise to substantially higher mutation rates in the natural mutator strains, this may not reflect the historical influence of this pair of alleles. In particular, an initial elevation of genomic mutation rate when the mutator alleles were first combined on a single genetic background could be dampened by the subsequent appearance of a modifier of mutation rate [25]. This scenario could manifest as an increase in genome-wide mutations in only those strains carrying the mutator combination (due to the historically higher mutation rate in those backgrounds) despite the present similar overall genomic mutation rates for both mutators and non-mutators.

To search for evidence of this pattern, we examined the number of derived mutations in each strain. We first aligned full genomes of the assayed strains to the genome of *S. paradoxus*, the closest extant relative of *S. cerevisiae*. Next we examined sites that were polymorphic in *S. cerevisiae* and for which we could determine the derived allele using the *S. paradoxus* sequence. We found that the number of derived alleles in natural mutator strains was not higher than the number in non-mutator strains (natural mutator mean 38,578; non-mutator mean 39,307). The same pattern held when we restricted our focus to the natural mutator strains and their two closest relatives at the whole-genome level (YJM 1775 and YJM1787) in order to focus on mutations that occurred along branches separating the natural mutators from their close relatives (natural mutator mean 30,103; non-mutator mean 30,316). Nevertheless, it is difficult to falsify the hypothesis that background modifiers of mutation rate arose after the incompatible mutator alleles were brought together in a single genetic background for at least two reasons. First, the mutation rate modifier allele(s) could have arisen shortly after the mutator alleles were brought together, resulting in a very small excess of mutations due to a short period of elevated mutation rate. Second, the complex population history of diverse yeast isolates, along with an unknown natural lifecycle (i.e., clonal growth vs. sexual reproduction), could obscure historical patterns of sequence divergence within and among lineages.

### No candidates for compensatory mutations modifying mutation rate

The lack of a strong effect of the mutator alleles on mutation rate in the clinical isolate backgrounds that we tested suggests a role for epistatic interactions with other loci. Previous work investigating this incompatibility utilized natural mutator alleles that contained additional SNP variants within *MLH1* and *PMS1* [42]. These results demonstrated that intragenic modifiers could modulate the mutation rate over a 20-fold range, although in all cases the mutation rate was still significantly elevated (at least 24X; [insert ref 42]). In contrast, our results suggest the presence of modifiers that greatly suppress the elevated mutation rate phenotype. Indeed, the slight elevation in mutation rate that we observed among natural mutator strains matches roughly with variation in mutation rate among different backgrounds with *compatible* allele combinations (up to 6-fold increase relative to S288c) [42].

We used sequence data [34] to search for possible intragenic modifiers in the set of strains we assayed. Demogines et al. [42] found two alleles – *PMS1*-F165C and *MLH1-*L271P – that modified mutation rate in strains with the incompatible mutator allele combination. The *PMS1*-F165C allele that increases mutation rate was not found among the strains we examined. However, the *MLH1*-L271P allele that decreases mutation rate 3- to 4-fold was polymorphic among the natural mutator strains. Strain YJM1785 (isogenic to YJM555) carries the reference T allele at position 595,697 on chromosome XIII, while the other three natural mutators carry the C allele that dampens the increased mutation rate due to the incompatible alleles. We observed a modestly lower mutation rate (~2-fold) among the natural mutators carrying this “protective” allele, in line with previous observations [42]. In addition to these variants, there were 18 SNP and indel variants in *MLH1* and 40 in *PMS1* that were polymorphic in the full panel of 100 Genomes strains we studied, of which 11 and 13 result in non-synonymous substitutions, respectively. Of these, only 2 SNPs in *MLH1* and 1 SNP in *PMS1* were polymorphic among the natural mutator strains. However, all had the same allelic pattern as the *MLH1*-L271P allele (YJM1785 carried one allele and the other natural mutators carried the other), preventing us from determining whether any of these additional non-synonymous polymorphisms also modulate mutation rate in the incompatible background.

We expanded our focus to the full genome to search for candidate loci that could contain mutation rate-lowering modifiers of the incompatible allele combination. First, we compared strains YJM320, S288c, and Sigma1278b (non-mutators that may not contain compensatory mutations) versus strains YJM541 and YJM554 (natural mutators that presumably *do* contain compensatory mutations). However, there were no polymorphic sites that had one allele shared by the mutators and another by the non-mutators. Next, we searched for sites where all natural mutators shared one allele but their closest relatives at the genomic level (YJM320 and YJM681) shared another allele. Unfortunately, we found over 1,000 sites that fit these criteria, precluding any realistic efforts to identify candidate modifiers. Overall, our inability to detect candidates may be because these strains are sufficiently divergent that a few large-effect compensatory mutations are obscured by the large amount of variation at other loci, or because there are many small-effect modifiers of mutation rate scattered throughout the genome. Moreover, these modifiers may have already been present in the genomic background on which the mutator alleles were first brought together. The complex machinery that is required to replicate DNA with high fidelity[55] makes it possible for many loci to influence the mutation rate. Given the challenges of making precise measurements of the mutation rate phenotype, uncovering modifiers of small effect using QTL approaches is not feasible.

### Conclusions

In this study, we characterized the mutation rate in strains derived from natural isolates carrying a pair of alleles in mismatch repair genes that were previously shown to lead to a much higher genomic mutation rate when combined in one background [41]. The existence of naturally-derived strains carrying this pair of alleles is unexpected given the prediction, from previous theoretical and experimental work, that natural populations of sexual microbes should not contain considerable frequencies of individuals carrying mutator alleles. We show that the mutation rate of laboratory strains engineered to carry these incompatible mutator alleles is high, as demonstrated previously [41], but that the mutator alleles only mildly elevate mutation rate in the strains derived from natural isolates. Thus, the effect of the incompatible alleles on mutation rate is modulated by genomic background.

The observation that all natural mutator strains identified by Strope et al. (2015) are derived from clinical isolates raises the possibility that the human body, an unusual environment for *S. cerevisiae*, poses unique selective pressures that could select for the mutator combination hitchhiking along with advantageous alleles during a period of clonal growth. Our data do not strongly support this contention given the relatively slight elevation of mutation rate in natural mutator strains. However, the haploid natural mutator strains we tested were derived from diploid clinical isolates that were likely highly heterozygous [40]. Thus, it remains a possibility, albeit unlikely, that within the population of *S. cerevisiae* present in a single patient, these alleles could be present at moderate frequencies along with other mutation rate modifiers such that rare offspring containing the mutator alleles and no suppressors could contribute adaptive mutations that are quickly detached from their high-mutation-rate background via recombination.

Given the possible link between mutators and opportunistic pathogens, the question of the effect of mutator alleles in pathogenic strains of a facultatively sexual microbe is of particular interest. Our results demonstrate that a cluster of clinical yeast strains with putative mutator alleles are in fact not mutators. More broadly, it is intriguing that hundreds of isolates of the facultatively sexual yeast *S. cerevisiae*—from around the globe and from various ecological niches— have been sequenced and the only isolates found to contain the incompatible mutator combination do not actually have a strongly elevated mutation rate. It is worth noting that mutator alleles are more likely to be maintained in facultatively sexual organisms than in obligately sexual organisms due to periods of clonal growth where such alleles cannot be decoupled from beneficial mutations. Given these observations, we believe our results provide rare and compelling evidence that supports evolutionary theory suggesting that sexual organisms are unlikely to harbor alleles that increase their genomic mutation rate.

## Data Availability

All data analyzed in this manuscript is available in Supplementary File 1.

## Author Contributions

HAM conceived of the study. BM and HAM carried out assays to estimate mutation rate. DAS and HAM analyzed the data. DAS conducted bioinformatic analyses of genomic variation. HAM and DAS wrote the paper. PMM provided strains and overall advice on the project. All authors gave final approval for publication.

## Acknowledgements

We thank John McCusker and Eric Alani for strains, and PD Sniegowski for helpful comments on the manuscript.

